# Mechanisms of alcohol influence on fear conditioning: a computational model

**DOI:** 10.1101/2023.12.30.573310

**Authors:** Adam Lonnberg, Marian L. Logrip, Alexey Kuznetsov

**Affiliations:** University of Evansville, Department of Mathematics, Indianapolis, Indiana, USA; Indiana University-Purdue University, Department of Psychology, Indianapolis, Indiana, USA; Indiana University-Purdue University, Department of Mathematical Sciences, Indianapolis, Indiana, USA

**Keywords:** computational model, electrophysiological activity, amygdala circuit, fear acquisition, fear extinction

## Abstract

A connection between stress-related illnesses and alcohol use disorders is extensively documented. Fear conditioning is a standard procedure used to study stress learning and links it to the activation of amygdala circuitry. However, the connection between the changes in amygdala circuit and function induced by alcohol and fear conditioning is not well established. We introduce a computational model to test the mechanistic relationship between amygdala functional and circuit adaptations during fear conditioning and the impact of acute vs. repeated alcohol exposure. In accordance with experiments, both acute and prior repeated alcohol decreases speed and robustness of fear extinction in our simulations. The model predicts that, first, the delay in fear extinction in alcohol is mostly induced by greater activation of the basolateral amygdala (BLA) after fear acquisition due to alcohol-induced modulation of synaptic weights. Second, both acute and prior repeated alcohol shifts the amygdala network away from the robust extinction regime by inhibiting the activity in the central amygdala (CeA). Third, our model predicts that fear memories formed in acute or after chronic alcohol are more connected to the context. Thus, the model suggests how circuit changes induced by alcohol may affect fear behaviors and provides a framework for investigating the involvement of multiple neuromodulators in this neuroadaptive process.

## Introduction

Alcohol use disorder (AUD) is a chronically condition characterized by loss of control over drinking as well as high propensity to relapse despite periods of abstinence (1). Stress is a predisposing factor for AUD (2) as well as other mental illnesses. The fact that past trauma increases the propensity to develop AUD, as well as being a primary trigger of relapse (3), implicates similar neurocircuitry underlie the maladaptive effects of stress and AUD. Specifically, neuroadaptations of stress circuitry contribute to the pathology of AUD (4) and are thought to be the point of intersection by which stress can promote the development of AUD. Among stress-related mental illnesses, PTSD in particular is characterized by rapid reacquisition of the fear response following a generally mild stressor, and AUD often potentiates these symptoms (5). Elucidating at the circuit level how stress and alcohol interact to alter neural adaptation, and the implication of those changes for behavior, is important for identifying methods to improve treatments for comorbid stress- and alcohol-related diseases.

Fear conditioning has long been studied to elucidate the impact of stress learning on neural circuit adaptation in rodents (6,7). Studies of fear conditioning focus on the initial learning of the fear behavior (acquisition) as well as on the processes by which fear-related behaviors may fade over repeated exposure to stress cues in the absence of the stressor (extinction). Extinction may be permanent or transient (8,9), dependent on the strength of physiological changes in the brain regions comprising the stress circuitry during formation of the extinction memory (10). Fear can be restored by a number of factors including the passage of time, change in context, or reappearance of the unconditioned stimulus (11). These studies provide a strong foundation for the behavioral research of the interaction between stress and alcohol effects in rodents and other model organisms.

The amygdala is well-known to be central to fear conditioning and extinction processes (12,13). The potentiation of connections between input nuclei of the amygdala and their respective inputs is central for the fear acquisition process. In particular, the increased strength of thalamic connections to the lateral amygdala and hippocampal projections to a subset of neurons in its basal nucleus are responsible for fear acquisition (14,15) (also see Methods). Fear extinction is new learning that strengthens another amygdala input, which suppresses the activity of fear-encoding neurons, rather than reversing previous fear learning-related changes (16,17). Specifically, cortical inputs to a subset of neurons in the basal nucleus are potentiated during extinction. Thus, the amygdala circuitry is the primary focus of research on the stress and alcohol interactions.

Stress and alcohol interactions are very complex and not understood in detail. Single or repeated fear conditioning sessions prior to alcohol exposure increase future alcohol intake (18,19). Several studies has shown impaired fear acquisition in acute alcohol (11,20,21). However, in several other studies acute alcohol has no effect on fear acquisition (22,23), or even causes its amplification at low alcohol doses (24). On the other hand, fear extinction has been consistently shown to slow down and weaken in acute alcohol (22,25).

Chronic alcohol exposure has variable effects on fear acquisition as well: facilitates (26) or causes no effect (27) in rodents or reduces in humans (28). It is further shown to facilitate pre-existing fear memories (29). Stephens shows reduction in fear acquisition for binge drinkers but sees generalization of fear in them. However, fear extinction has been shown to consistently become weaker after chronic alcohol exposure (26,27,30). In this paper, we aim to reproduce the impaired fear extinction by incorporating known alterations produced by alcohol in the amygdala circuitry in a computational model.

We connect the alterations produced by alcohol at the circuit and behavioral levels in a computational model. Modeling can bridge the gaps between in vivo and in vitro techniques used to investigate behavior and neural activity changes. Specifically, we analyze how altered amygdala connectivity changes behavioral responses in fear acquisition and extinction. We simulate two conditions: first, acute alcohol and, second, prior repeated alcohol, Thus, the aim of this study is to establish mechanistic relationships between amygdala function and structural adaptation during fear learning and the impact of acute and chronic alcohol. This model will provide a framework for investigating putative modulators of amygdala circuit activity that can inform future behavioral and translational research.

## Methods

### Amygdala Structure

#### Basolateral amygdala

The basolateral amygdala (BLA) is a crucial component of the brain’s limbic system, playing a pivotal role in the processing of emotions such as fear and anxiety (31). The BLA sends glutamatergic projections to the central amygdala and other downstream targets. The BLA was separated into lateral (LA) and basal amygdala (BA) according to their distinct inputs and functional differences (Fig. 1). The LA, which is crucial to fear acquisition, receives thalamic and cortical inputs (REF). The BA contains two functionally opposing groups of projection neurons, One of the groups, referred to as fear neurons (BAf), are activated in response to aversive stimuli and are essential for the expression of conditioned fear responses (Fig. 1) (32). These neurons receive inputs from the LA and hippocampus (14,15,17), and excite downstream pro-fear neurons in the central amygdala (CeAOn) as well as other brain regions involved in the expression of fear behaviors (33). Activation of these BLA fear neurons is associated with increased heart rate, blood pressure, and freezing behavior, all of which are characteristic physiological responses to fear (34).

**Figure 1:**
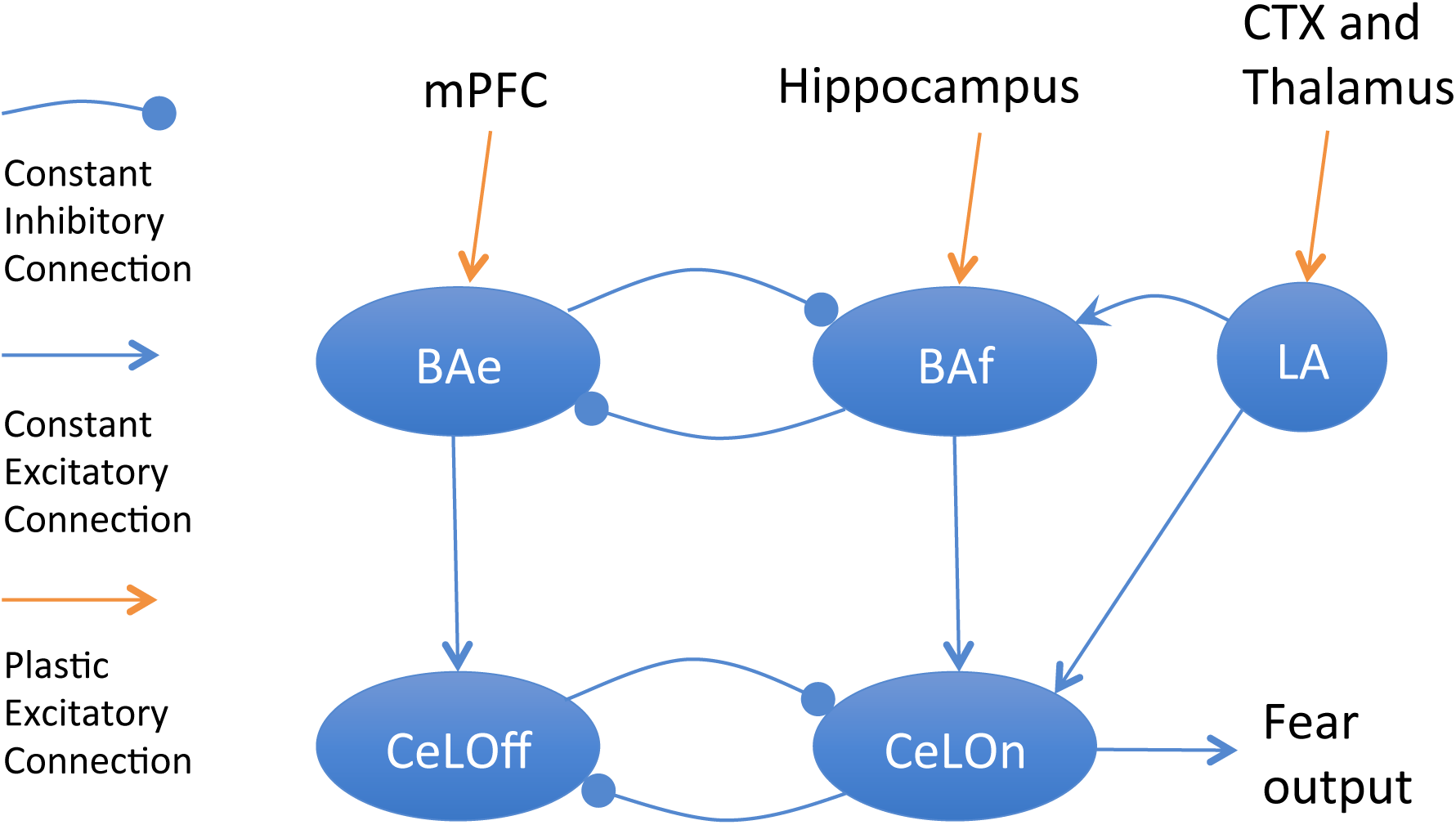
Diagram of the amygdala circuitry implemented in the model.

Conversely, another group of projection neurons in the BA, referred to as extinction neurons (BAe), is involved in the inhibition of fear responses (35). These neurons receive input from the medial prefrontal cortex (mPFC) (14). During extinction learning, where a previously conditioned fear response is inhibited by repeated exposure to the conditioned stimulus without the aversive outcome, there is an increase in the activity of the extinction neurons, and a decrease in the activity of the fear neurons (35). This shift in neuronal activity is associated with a reduction in fear expression and an increase in approach behaviors towards the conditioned stimulus (36). Activation of these neurons is associated with a reduction in heart rate, blood pressure, and freezing behavior, and an increase in approach behaviors towards positive stimuli (37).

A growing body of evidence suggests that the BA fear and extinction neurons are mutually inhibiting (Fig. 1) (31,38–40). The antagonistic relationship between these two groups of neurons in the BLA is essential for the appropriate regulation of emotional responses.

Extensive research has shed light on the roles of inhibitory interneurons within the BLA and their involvement in shaping emotional processing, particularly in aspects related to fear and safety learning (41–45). These interneurons play an instrumental role in controlling the input-output dynamics of the BLA by modulating the excitability and timing of projection neuron activity (44,46). While their contribution in gating inputs to the BLA is relatively well-understood (33,47), the exact mechanisms by which they might modulate mutual inhibition between neurons associated with fear and those linked with safety remain ambiguous (39,48). This uncertainty underscores the need for further exploration and, potentially, the development of a computational model to provide clarity. Therefore, in our present model, we omit details of these connections and code the interaction between the BA neural groups as direct mutual inhibition (Fig. 1).

#### Central amygdala

The circuitry of the central amygdala (CeA) is also quite complex and underwent extensive investigation (for review see (15)), although many details are still uncertain. Therefore, we reduce the circuitry to functionally minimal. As well as the BLA, the CeA is divided into pro-fear (CeAOn) and pro-extinction (CeAOn) neurons (15) (REF). However, these neurons are all inhibitory and directly inhibit one another (REF). The pro-fear neurons in the medial part of the CeA project to downstream targets and induce fear-related behaviors (15). This provides another layer of inhibition between the fear and extinction pathways (Fig. XX). We group together the pro-fear neurons from medial and lateral parts of the CeA in the group denoted as CeAOn as their function and inputs are similar.

BLA directly projects to the central amygdala (49) (10.1016/j.neuron.2017.02.034). Specifically, LA neurons connect to the pro-fear neurons in the medial part of CeA and BA fear neurons – to the lateral. As we united these pro-fear neurons into the same group, CeAOn, both inputs excite this group. On the other hand, the extinction neurons from BA, BAe, project to the pro-extinction neurons in the central amygdala, CeAOff (10.1016/j.neuron.2017.02.034). Therefore, the BLA to CeA connections comprise mutually inhibiting pro-fear and pro-extinction pathways (Fig. XX). The output of the model is the pro-fear CeAOn grout since these neurons from the medial part of the CeA project to downstream targets (REF).

The amygdala circuit contains other groups of neurons that mediate its signal transduction. In particular, two clusters of intercalated (ITC) neurons dorsolateral (ITCd) and ventralateral (ITCv) mediate BLA to CeA pathway. However, we have not included them in this model as they repeat the connections that the model accounts for. Specifically, ITCd neurons connect LA to CeAOff neurons (49), which provides inhibition of the latter in a similar way as connection from LA to CeAOn neurons as CeAOn neurons inhibit CeAOff neurons in the model. Similarly, ITCv neurons are excited by BA extinction neurons and provide inhibition of CeAOn neurons (49) in a similar way as BA fear neurons reduce their excitation due to their inhibition by the BA extinction neurons. In some circumstances, these two additional pathways may act differentially from those they are functionally similar to, but this requires their differential inputs. Therefore, we merge the similar pathways and augment the strength of the LA to CeAOff and BAf to CeAOn bisynaptic connections to account for the reduction.

#### Firing rates of neural groups

For each neural group *i* in the model, the firing rate is determined by the following equation common for all neuron populations (50):

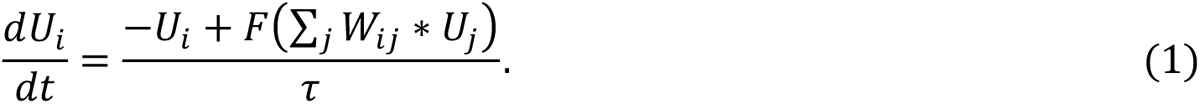

Here, *U_i_* is the firing rate of neural group *i*, *W_ij_* is the synaptic weight between group *i* and group *j*, τ is the time constant set to 0.05 sec (51), and *F* is the sigmoidal function

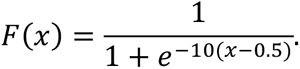

We model a total of five neuronal populations using these equations (Fig. 1). The structure is the same as in earlier modeling (51).

### Synaptic Plasticity

Fear acquisition and extinction are linked to plasticity of synaptic connections within the amygdala and its inputs (52–54). Among them, potentiation of synaptic inputs that transmit stimuli and context-related information to amygdala is shown to be key in modulation of fear responsiveness (15,53–60). Therefore, we assume connection strength of the three inputs: from the thalamus to the LA, from hippocampus to BAf, and from mPFC to BAe, all to be plastic. To focus on the contribution of plasticity of these three inputs, all other connections are given constant synaptic weights. An important simplification achieved by this assumption is that task-dependent plasticity and alcohol-dependent plasticity affect different synapses in the model. This simplification is necessary to avoid extreme complexity due to mutual interdependence of plasticity processes. The plastic synaptic weights of the inputs to the LA and the BAf implement a variation of the Rescorla-Wagner (Hebbian type) learning rule (61):

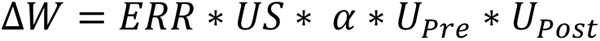

Where α is a plasticity rate constant, the US is the adverse unconditioned stimulus that engenders learning and the *U_pre_* and *U_post_* are the respective firing rates of the pre-and post-synaptic neurons. The prediction error is calculated as

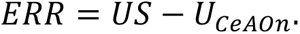

Note that the accepted notation is to register an adverse unexpected stimulus as positive prediction error, whereas a negative error is an unexpected omission of the stimulus (17).

For the synaptic weight projecting to BAe, the following was used:

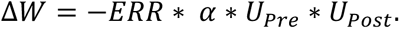

In other words, only in the presence of a shock (US=1), the inputs to the fear pathway are potentiated. In the absence of a shock (US=0), the inputs to the extinction pathway are potentiated until the prediction error vanishes as *U_ceLOn_* → 0. Using these rules, a negative prediction error leads to inhibition of the fear pathway not by a decrease in a synaptic weight, but by potentiation of an alternative pathway (17). This is justified given that little evidence shows synaptic depression in either fear acquisition or extinction, and that it is widely accepted that the activation of alternate pathways is key in fear extinction (61,62).

### Behavioral Protocol

The learning protocol involves two periods: fear acquisition and fear extinction. The acquisition period consists of first 25 trials and the extinction period extends to additional 35 trials. Each trial has three phases (14,51) (Fig. 2). The first phase is the presentation of a context and the conditioning stimulus (CS) in a context specific for it. In the model, the CS and the context are represented as continuous signals from the thalamus to the LA and the hippocampus to the BA respectively (17,63). After this phase, the network’s prediction of an aversive unconditioned stimulus (US)—that is, the activity of *CeLOn*—is read out. The second phase then begins, during which all signals from the previous phase continue, and a US is added in acquisition. In extinction, no US is presented. At the end of phase two, all synaptic weights are updated. Finally, during the third phase, no input is presented to the system allowing it to reset. The first phase of the next trial then begins.

**Figure 2:**
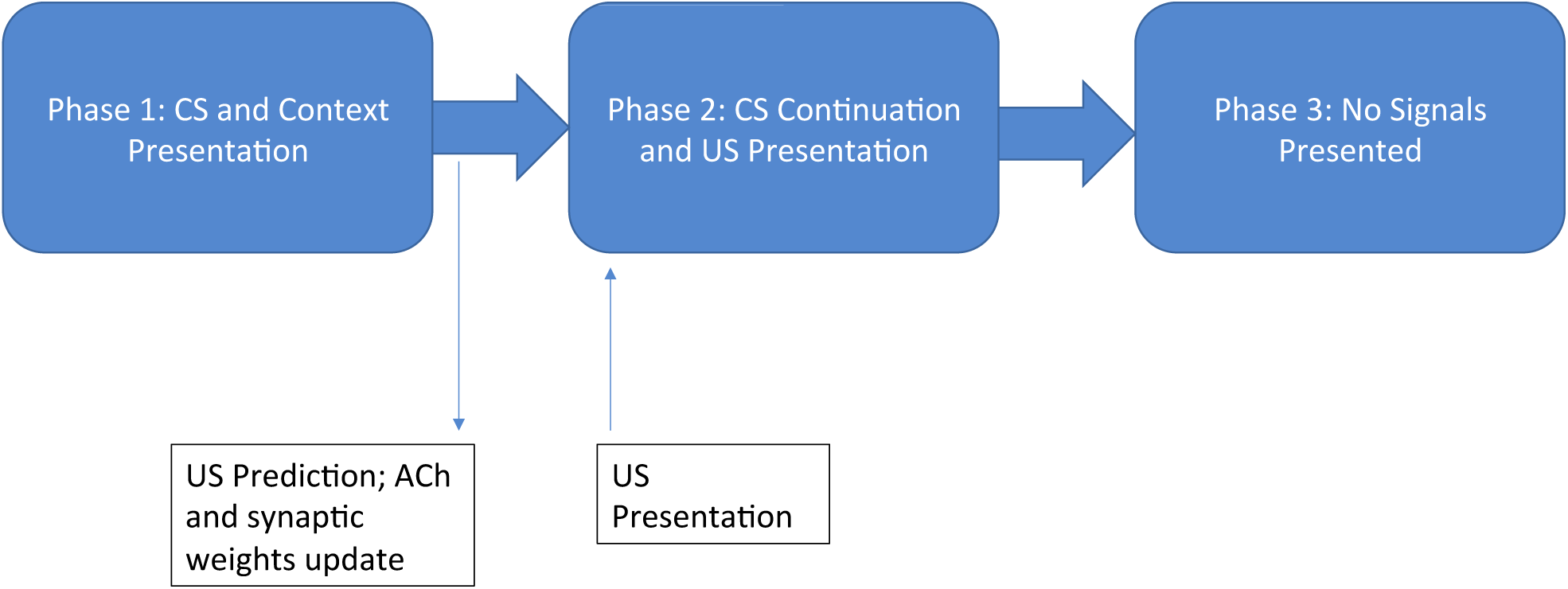
The learning protocol used in all trials in the model.

Extinction also takes place in a different context, as in the experiments (14) and prior modeling (51,63). Thus, the hippocampal input to the BAf is turned off in the model because the new context is not associated with fear. By contrast, an input from mPFC to BAe is activated during extinction (14,51), and synaptic connections from mPFC to BAe start to grow according to the plasticity rules described above.

### Calibrations and Simulations

Our model is coded in, and all results were generated by Python. We used Euler’s method, which is sufficient for solving this system given its strong convergence to steady states. All nuclei include additive normally distributed noise. A step size of 2 milliseconds is used, and each trial described above consists of 500 timesteps. We average the number of trials needed for fear acquisition and extinction over 100 model instances and calculate their standard deviations.

To calibrate the model (Table 1), we combine functional criteria of fear acquisition and extinction with available electrophysiological data on synaptic currents and firing rates of each neural group. First, we use current-frequency relation for CeA neurons (64) to calibrate the input response function *F*(*x*): a unit of input in the model represents ∼ 100pA current. We generalize this scaling to all neurons in the model to compensate for sparse data. For CeA neurons (including CeA On and Off groups) the EPSP and IPSP amplitudes were measured to be approximately 65 and 150pA respectively (26,27). While these data are a good initial approximation for the model, it lacks the information about the convergence of the inputs, which affects the activation of these currents. This gives us some freedom to calibrate the synaptic strengths to reproduce the behavioral data on fear responses. For this reason, we assign a greater weight for the connection from BAe to CeAOff to robustly reproduce fear extinction (*W_BAe-CeA_*; Table 1). The IPSP amplitude in LA neurons were up to 100pA (65,66), and, thus the inhibitory synaptic weight within LA was calibrated to be 1 in the model. Finally, the weight of LA to BAf connection determines the residual BAf response to CS after the animal is switched to another context (e.g. extinction) (15). The strength of this connection was not measured directly, and we use this functional criterion.

**Table 1:**
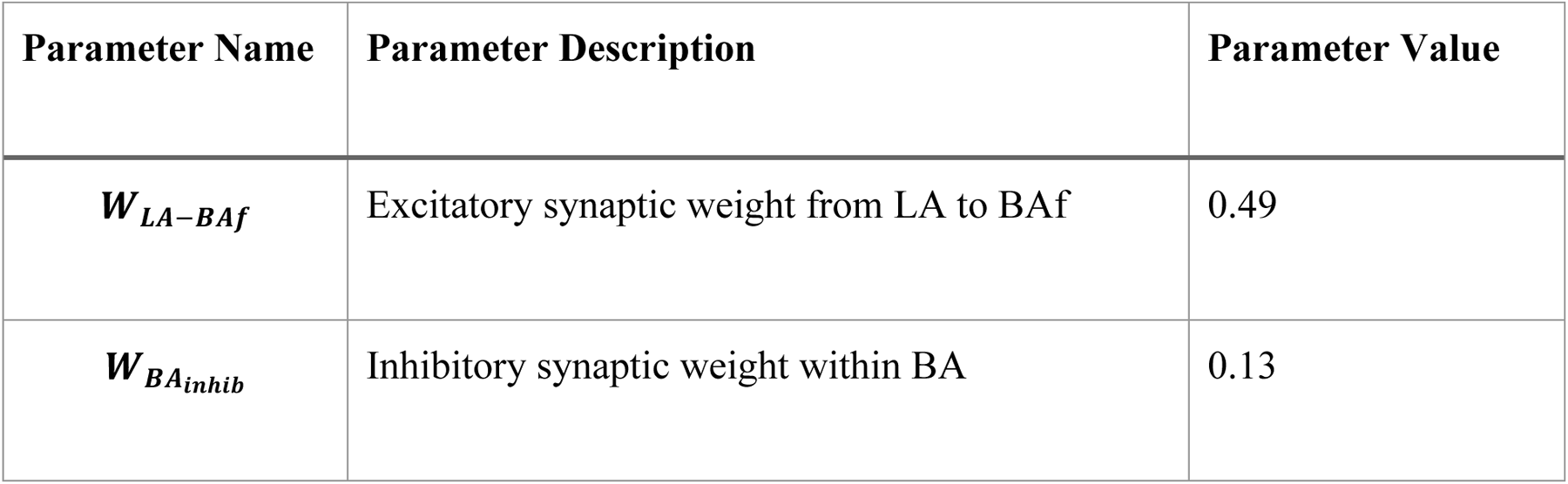
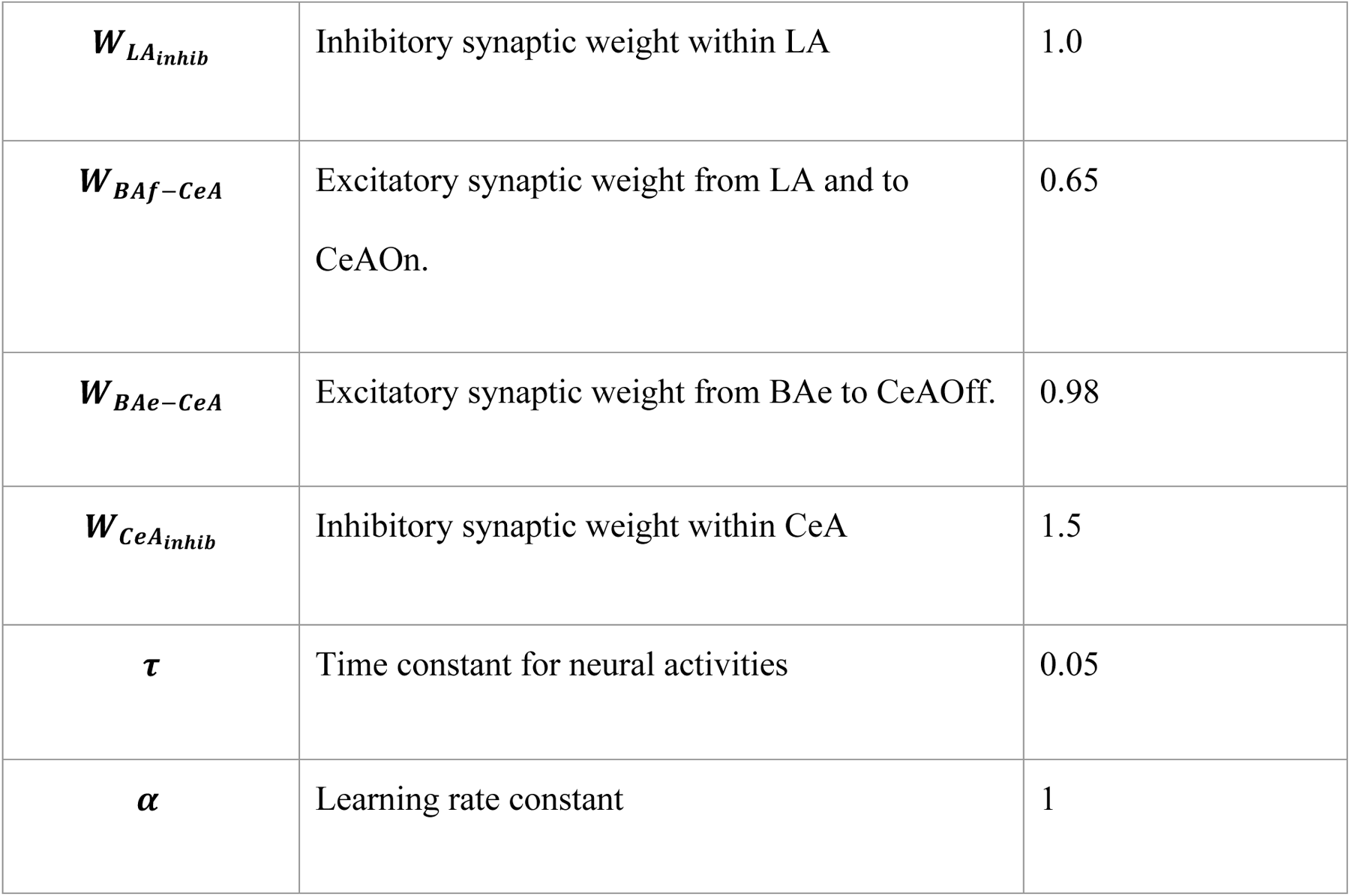
Model parameters. Values are calibrated as described in Methods.

All activity levels were normalized to be between 0 and 1 for this model. Functionally, the above calibration resulted in full activation of CeAOn (fear expression) at a partial activation of BAf and LA. Thus, this calibration ensures robustness of fear learning since fear expression requires only partial activation of the basal amygdala fear pathway.

Next, we test functional robustness of the sub-system responsible for fear extinction: the BAe and CeAOff (Fig. 1). Fear extinction is achieved by inhibition of CeAOn and BAf by activation of mPFC synaptic input. Analysis of the topology of the model allows us to give conditions on parameters under which extinction is possible. Namely, the sum of the inhibitory and excitatory inputs to CeAOn must be below the threshold for the response function F(x):

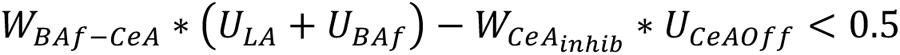

Thus, extinction occurs if 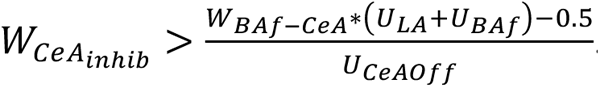 . The value of this parameter in Table 1 is well within this range, which ensures robust fear extinction. Note that we used the same connection weight *W_BAf-CeA_* for both LA and BAf inputs to CeAOn because the data do not differentiate the neural groups. Lastly, our calibration of mutual inhibition between BAe and BAf (67) allow for fear- and extinction-responsive neurons in BA coactivate, as shown in experiments (14). Thus, we assign a much lower value of 0.13 for the inhibitory synaptic weight within BA.

### Alcohol-Induced Modulations

Alcohol modulates the weights of the connections between various subdivisions of the amygdala as well as its afferents. Acute alcohol leads to increased inhibition in the BLA and CeA. Specifically, GABA transmission was increased by ∼50% (66,68–70). Simultaneously, acute alcohol reduces glutamatergic transmission in CeA by ∼20% (71). Accordingly, we modulate the synaptic weights to reflect these changes in the model (Table 2)

**Table 2:** Parameter modulation by acute alcohol. All other parameters stay the same.

| Parameter | Modulation | Literature |
| --- | --- | --- |
| $W_{BA_{inhib}} = 0.25$ | Increased by ~50% | (66,68–70) |
| $W_{LA_{inhib}} = 1.5$ | Increased by ~50% | (66,68–70) |
| $W_{BAf-CeA} = 0.4$ | Decreased by ~20% | (71) |
| $W_{BAe-CeA} = 0.6$ | Decreased by ~20% | (71) |
| $W_{CeA_{inhib}} = 2.5$ | Increased by ~50% | (66,68–70) |

In chronic alcohol, the most prominent change observed is a four-fold increase in the baseline GABA levels in the CeA (71). However, this research shows a much more modest increase in the IPSC amplitude (∼60%), and, accordingly, we strengthen the inhibitory interaction in CeA by this amount (Table 3). The rest of the increased GABA levels are hypothesized to increase a tonic inhibitory component in the CeA (72). Thus, we introduce constant parameters (drive: *Dr_CeA_* = −0.5) that reduce the input current in Eq. (1) and inhibit CeA neuron firing. There are also reports of increased glutamate transmission in the BLA in the chronic alcohol condition (73). In the CeA, although, such reports are somewhat inconsistent (74), except for a response to an alcohol challenge (75). To account for this increase, we strengthen the input from LA to BAf neurons (Table 3). The other three excitatory connections in the BLA are inputs, and their plasticity is defined by the conditioning protocol (see above). We will see that, indeed, they undergo greater potentiation in the chronic alcohol conditions (see Results).

**Table 3:** Parameter modulation by chronic alcohol. All other parameters stay the same.

| Parameter | Modulation | Literature |
| --- | --- | --- |
| $W_{CeA_{inhib}} = 2.5$ | Increased by ~60% | (71) |
| $Dr_{CeA} = -0.5$ | Increased tonic GABA levels (4-fold) and IPSC frequency (0.22 to 0.72 Hz) | (72) |
| $W_{LA-BAf} = 0.6$ | Increased by ~20% | (73) |

## Results

### Fear Acquisition

The first 25 trials of the simulations tested fear acquisition. During these trials, the model network receives a CS input activating the LA and a context input activating the BAf. The inputs are maintained until the presentation of a US (shock) in the second phase of each trial. The US is represented in the calculation of the prediction error (see Methods).

Fig. 3 shows trial-by-trial dynamics of the averaged activity of all amygdala neuron types we model (A,B), and plastic synaptic weights (C). The last panel (D) shows exemplar neural activity within first several trials. Acquisition trials are to the left of the dashed vertical line and numbered 1-25. The activity of each neural group is averaged across phases 1 and 2 of each trial (see Fig. 2). As the figure shows, the activity of all neural groups is low at the beginning of acquisition process. The initial growth of activity is very slow followed by a sudden increase in activation of LA, BAf, and CeAOn groups. The process takes on average 6.26 trials (std 0.44), after which activity remains at approximately the same level. The behavioral readout of the model is activation of the CeAOn neurons, which shows fear response to the CS presentation. This completes the fear acquisition process.

**Figure 3:**
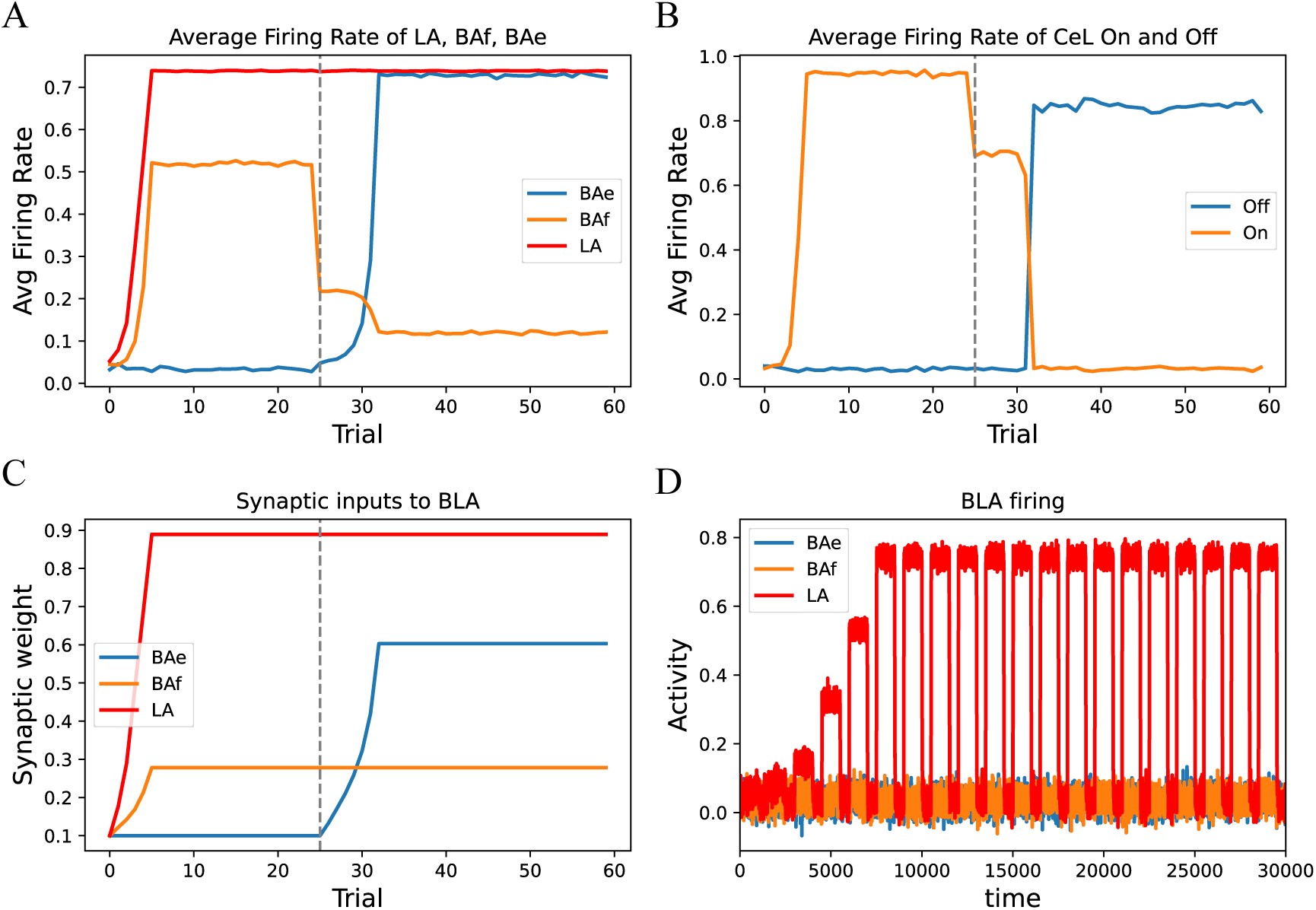
Dynamics of amygdala activity and synaptic inputs during fear acquisition and extinction. Acquisition trials are 0 to 24. Extinction trials are 25 to 59. The vertical dashed line separates these two periods. (A) Average activity levels of LA BAf and BAe. (B) Average activity levels of CeA On and Off . (C) Weights of synaptic inputs to LA, BAf and BAe. (D) Exemplary time dependence of the activity levels throughout the first 20 trials for the BLA neural groups. The averaging in (A) is done over the first two phases of each trial, whereas the third phase is an intertrial interval, during which the activity is reduced to the background levels.

The mechanism for fear acquisition in the model is potentiation of the synaptic weights from the sensory areas and hippocampus to the LA and BAf respectively. Their dynamics are shown in Fig. 3C. Potentiation of these two projections is due to a positive prediction error: the difference between the punishing US presented on phase 2 and the lack of CeAOn activation. However, the initial growths of the synaptic weights are very gradual. This is due to Hebbian plasticity rule (see Methods), which makes the changes in the synaptic weights dependent on the activity levels of both pre- and post-synaptic neurons. Thus, the low initial activity of LA and BAf impedes fast potentiation of the synapses. The prediction following from this mechanism is that fear acquisition can be accelerated by directly priming the activity of the LA and/or the BAf neurons.

### Robust Fear Extinction

Fear extinction was tested beginning in trial 26 until the end of the simulation. During these trials, the CS continues to activate the LA as it did during the acquisition trials. The context, however, changes. A set of hippocampal neurons responsive to the new context is not associated with fear and, thus, not activating the BAf. Furthermore, as extinction develops, projections from a set of mPFC neurons responsive to this context potentiate and activate the BAe (14). In the model, therefore, we silence the input from the hippocampus to the BAf and activate mPFC neurons that provide weak initial stimulation of the BAe. Additionally, no US is presented during extinction trials (i.e. no shock, which is incongruent with the presence of fear response).

Fig. 3 shows that at the beginning of extinction there is a substantial immediate drop in the activation of BAf (A). This occurs because of no excitation of BAf by the hippocampus due to the change in context. On the other hand, there is a very modest drop in the activation of CeAOn (Fig. 3B) due to persistent excitation from the LA (Fig. 3A). Therefore, the model reproduces initial pre-learned fear responses to the CS presentation regardless of the context. The persistent activity of the LA reflects its direct activation by the CS.

The immediate drop induced by the context change is followed by a gradual further decrease in the activity of BAf and CeAOn as well as a gradual increase in the activity of BAe. This behavior continues for 7 trials, after which there is a sharp increase in BAe and CeAOff activity.

Simultaneously, the activity of BAf and CeAOn sharply decreases. After this occurs in trial 33, there is little change in the activities of any of the system’s neural groups. The activity of LA remains constant throughout this process simply reflecting the same CS presentation in all trials. As the activity of CeAOn remains near zero, we register fear extinction. On average, it takes 8.54 trials (std 0.57).

The mechanism for extinction is the potentiation of the synaptic weight between the mPFC and the BAe. Fig. 3C shows the slow increase in this weight following trial 25, which is gradual at first due to the low activity of BAe, the postsynaptic neural population. The increase in the rate of potentiation corresponds with the increase in activity levels of the BAe. It is important to note that extinction does not occur by the depression of any of the synaptic pathways potentiated during acquisition (Fig. 3C red and orange). Rather, it occurs by the strengthening of an alternative pathway, which inhibits fear-responsive neural groups within the amygdala circuitry. Thus, mutual inhibition within amygdala determines the capacity for extinction.

### Acute Alcohol Impedes Fear Extinction by differentially affecting BLA and CeA activity

As mentioned in Methods, acute alcohol makes the excitatory inputs to CeA weaker and inhibition within amygdala, both in LA and CA, stronger. While the direction of the changes is clear, the magnitude is hard to calibrate due to insufficient experimental data. In particular, changes in activity of the amygdala recorded in vivo, whereas changes in synaptic function, such as transmitter release or receptor activation, are measured in vitro. To close this gap, we conducted parametric analysis by varying the parameters in a wide range and analyzing the changes in the activity simulated by the model. The analysis shows that the above parameter modulations exert robust activity changes in a wide range of physiologically relevant values. Based on this parametric analysis, we set the magnitude of parameter modulations that produced typical activity responses. In particular, Fig. 4 depicts the results obtained when parameters modulated by acute alcohol were set to the values in Table 2.

**Figure 4:**
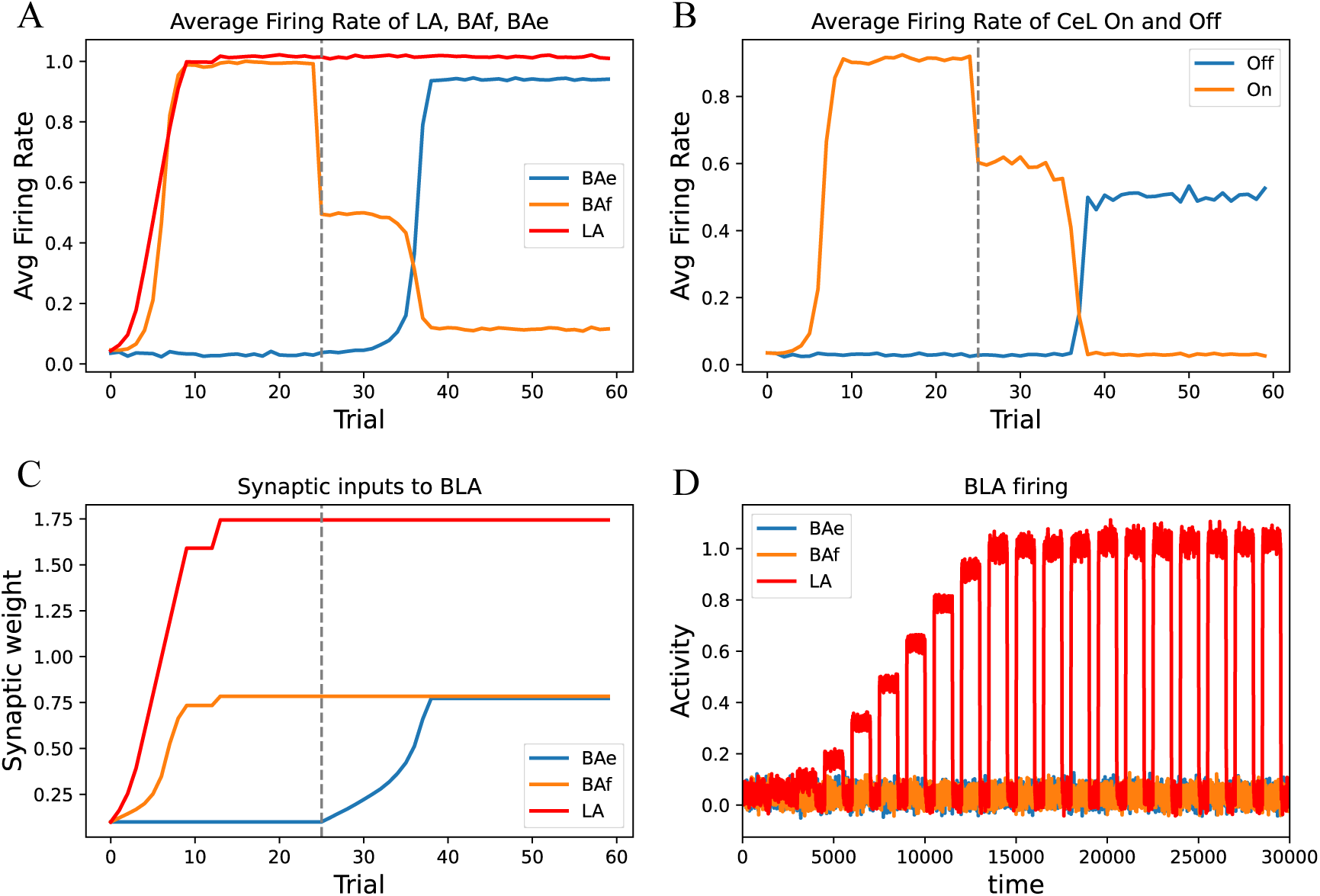
Effects of acute alcohol on amygdala activity during fear acquisition and extinction. Notation is the same throughout the panels as in Fig. 3. Parameter values changed from the control are in Table 2

Fig. 4 indicates that acquisition completes after an average of 10.43 trials (std 0.59), which is 3 trials longer than in the control conditions (compare with Fig. 3). This is directly caused by the reduction in the BLA to CeA connection strength. Since, as before, conditioning is contingent on the sufficient excitation of the CeAOn neurons, stronger excitation of their inputs is required to compensate for the reduced synaptic weight. Thus, the activation of LA and BAf reach greater values (Fig. 4A) due to greater potentiation of the corresponding connections (Fig. 4C). The additional potentiation requires the greater number of trials. Furthermore, the LA activation reaches its maximum and the additional increase of the overall input to CeAOn is achieved by further activation of the BAf neurons (Fig. 4A vs 3A). Therefore, our simulations predict that acute alcohol increases the contribution of BAf to fear conditioning.

Extinction takes far longer, increasing from 8.54 trials in the control condition to the average of 13.58 trials (std 0.49) in the acute alcohol condition (Fig. 4 trials 25 to 38). Thus, the model reproduces delayed extinction in acute alcohol (22,25). This delayed extinction, in large part, is a consequence of the stronger activation of the LA and BAf during fear acquisition. This stronger activation of the fear pathway constitutes a larger barrier to overcome in order to inhibit the fear response. Not only the activity of the BAe must now reach a greater level for extinction, but its activation is also impeded by inhibition from the BAf neurons. This slows down potentiation of the BAe input throughout extinction since the potentiation rate explicitly depends on the BAe activation (see Methods).

The increase in inhibition strength between BAf and BAe in acute alcohol further impedes fear extinction. The mechanism is the same: the stronger inhibition amplifies the influence of the fear response in the BAf on the BAe activity and does not allow it to grow.

Interestingly, the increased contribution of BAf to fear conditioning has only a minimal effect on fear extinction. Indeed, given that the BAf activation drops as the agent is shifted from acquisition to extinction context (Figs. 3A and 4A orange, trial 24 to 25), this makes extinction easier (and thus faster). Specifically, the activation level of the CeAOn neurons decreases following the context shift by ∼25% in control conditions and ∼35% in acute alcohol (Figs. 3B and 4B orange). Although this should enhance extinction, it becomes slower in acute alcohol.

Our simulations show that the increased inhibition in the CeA in acute alcohol does not significantly affect the behavior during conditioning but contributes to the reduction of the CeA activation levels (Fig. 4B).

Overall, the model predicts that fear acquisition and extinction in acute alcohol lead to greater activation of the corresponding neural groups in the BLA, but lower activity in the CeA. Most significantly, in acute alcohol, the activity of the CeAOff neurons after fear extinction is reduced most (∼40%) compared to that in control conditions (Fig. 4B blue).

### Chronic Alcohol Reduces Reliability of Fear Extinction by Inhibiting Activity of CeA

As we state in Methods, prior repeated alcohol exposure increases the inhibition in the CeA and excitation of the BAf by LA (see Methods and Table 3). Fig. 5 shows that the number of trials to fear acquisition is slightly greater after repeated alcohol exposure (8.12 vs. 6.26; std 0.32). However, activation of the fear pathway is substantially changed: LA activation achieves a somewhat greater maximum value, and BAf activation is greatly increased. These changes are similar to those in acute alcohol, but this causes a higher input to the CeAOn neurons as the CeA synaptic inputs remain the same in chronic alcohol. The higher CeAOn excitation is required for fear conditioning in chronic alcohol because it counteracts the amplified tonic inhibition in CeA. Thus, the modulations to the amygdala produced by chronic alcohol, however different from acute alcohol, produce very similar changes in the activity levels (Fig 4A,B and 5A,B).

**Figure 5:**
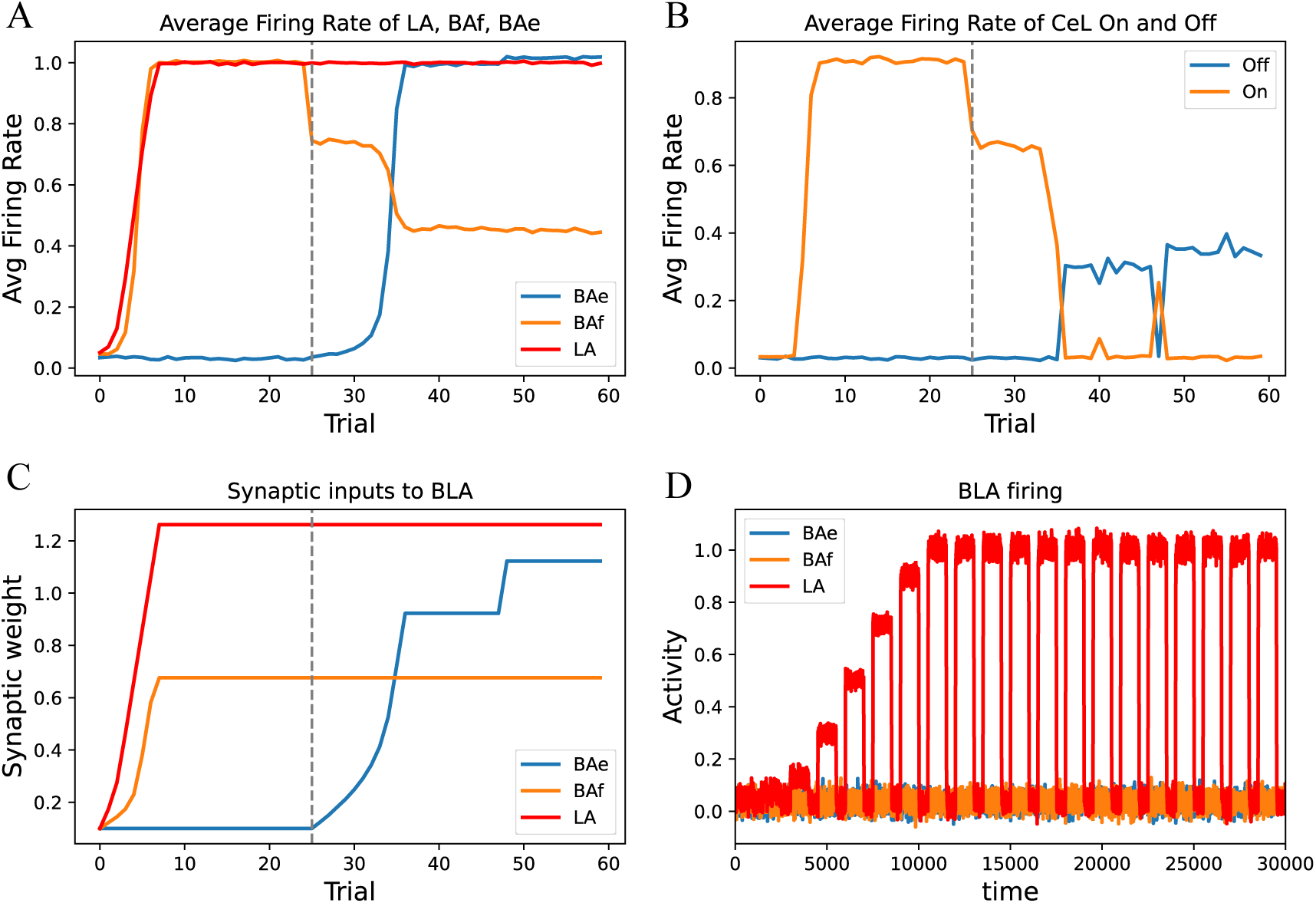
Effects of chronic alcohol on amygdala activity during fear acquisition and extinction. The notation is the same as in Fig. 3. Parameter values changed from the control are in Table 3.

Fear extinction trials start with substantially greater residual activation of both the LA and the BAf (Fig. 5A trial 25). Because of that, extinction is achieved later than in control conditions and takes on average 12.33 trials (std 0.53). Thus, our simulations reproduce impeded fear extinction after chronic alcohol (26,27,30). Furthermore, the activation levels of the CeAOff neurons after extinction, although sufficient for the suppression of fear response, remain low. This change parallels what we have shown above for acute alcohol but is amplified. Indeed, the CeAOff activity after extinction is down additional 20% compared to acute alcohol case (Fig 4B vs. 5B, blue). This activation level is so low that noise added to our simulations can switch the CeA and induce a fear response (Fig. 5B trial 47). In the simulation, the system has the capacity to further strengthen the extinction pathway (Fig. 5C), but the activation of the CeAOn neurons remains low (Fig. 5B trials 48-60). Therefore, fear extinction is much less reliable after chronic alcohol due to inhibited activity of the CeA neurons.

## Discussion

### Acute and chronic alcohol delay fear extinction

Acute (11,20–24) and chronic (26–28) alcohol has variable effect on fear acquisition. In our simulations, alcohol significantly increases the number of trials required for acquisition in both acute and chronic conditions (10.43 and 8.12 respectively vs. 6.26), and this trend replicates several studies (11,20,22,28). Our model suggests that the variability of the acquisition time may come from variable background levels of activity in the BLA or in the brain regions projecting to it prior to learning. The learning process involves Hebbian synaptic plasticity and is primed by the activity of both pre- and post-synaptic neurons (61). A greater initial activity level speeds up fear acquisition, whereas lower initial activity delays acquisition. Such priming can be done optogenetically, or by drugs such as alcohol. If these levels prior to learning were increased by alcohol, the acquisition time would be decreased to the values observed in the control conditions. However, for example, hippocampal activity is reduced by acute alcohol (76,77), and this should further slow down context-dependent learning. In our simulations, we assume the same initial activity levels in all conditions to avoid variations in the acquisition time. However, our modeling suggests that fear acquisition time may be highly variable as it depends on the background activity levels in the BLA and its input structures as well as the specifics of the task.

On the other hand, fear extinction is delayed more consistently by both acute (22,25) and chronic (26,27,30) alcohol. Our model successfully reproduces this result (13.58 and 12.33 trials to extinction respectively in acute and chronic alcohol vs. 8.54 in control). What mechanisms control the onset of fear extinction? The simulations show that the number of trials required for fear extinction is largely independent of that required for conditioning. For example, the above predicted variability in fear acquisition timing will not affect extinction. By contrast, the final levels of activity in the fear pathway reached after acquisition affect extinction. In particular, in both acute and chronic alcohol, the LA and BAf are activated stronger than in the control conditions, and, therefore, BAe receives stronger inhibition from BAf at the beginning of extinction. Accordingly, the longest extinction delay was obtained in the acute alcohol simulations, where BAf activity was greater by nearly a factor of two compared to the control case (Fig. 4A vs 3A, orange), and the BA inhibition weight was increased as well. Second, the extinction delay is affected by the background firing rate of the mPFC input. Indeed, by the same Hebbian rule, this extinction delay depends on the background firing rate of the postsynaptic BAe and the presynaptic mPFC neurons. In our simulations, we assume the same mPFC activity levels in all conditions to focus on the influence of the amygdala structure on fear conditioning. Therefore, the delay in fear extinction in alcohol is mostly induced by greater activation of the BLA after fear acquisition due to alcohol-induced modulation of synaptic weights.

### Fear Extinction is Less Robust in Acute and Chronic Alcohol

In our simulations both acute and prior repeated alcohol lead to greater activation of the BLA fear pathway after fear acquisition. To counteract this, in extinction, a greater activation of the BAe neurons is required as well. However, this does not lead to a greater activation of the CeA neurons in the simulations. To the contrary, all CeA neural groups are less active in alcohol. Indeed, the greater activation of the BLA is a compensatory mechanism that counteracts a greater inhibition and/or lower excitation in the CeA. The decrease in activity of the CeA is most significant for the CeAOff neurons in extinction for the chronic alcohol case (Fig. 5B, blue), where their activity is reduced by ∼60% compared to the control conditions (Fig. 3B, blue). As a result, the activation of the extinction pathway through CeA is insufficient to reliably oppose fluctuations in the fear pathway and, therefore, exclude fear responses. Thus, the model predicts that acute and especially prior repeated alcohol shifts the amygdala network away from the robust extinction regime.

### Alcohol Exposure Changes Context and Stimulus Contributions to Fear Conditioning

The aim of this study was to model the mechanistic relationships between amygdala function and structural adaptation in fear learning and the impact of acute and repeated alcohol on this process. In the model, two pathways mediate fear conditioning: the LA and BAf projections to CeA. These two pathways may contribute differently depending on experimental conditions. For example, BAf is activated more than LA in fear acquisition simulations when acute alcohol is present as well as after prior alcohol exposure, whereas LA contribution is greater in control conditions. The amplified BAf activity is caused by potentiation of hippocampal inputs and, thus, increases the contribution of context to the fear memory. Therefore, our model predicts that fear memories formed in acute or after chronic alcohol are more connected to the context.

The contributions of the conditioned stimuli and context during fear acquisition can be further manipulated by priming the activity in one of the regions that provide their excitation: e.g. thalamic areas for the LA and the hippocampus for the BAf. In particular, acute alcohol has been shown to drastically decrease hippocampal activity (76,77), which would compensate for the greater potentiation of the hippocampal input to the BAf neurons. The current version of the model does not account for these modulations but provides a framework for future expansion of the model’s predictive function. In particular, the model predicts that an attempt to restore hippocampal activity in alcohol to rescue contextual learning will exaggerate the context contribution to fear acquisition.

### The Model as a Framework for Elucidating the Impact of Modulators

A central purpose for development of this model is the generation of a framework that may be used to examine the impact of a variety of modulatory influences on fear learning. Here we limited the factors incorporated to first build a basic model. There are multiple neuromodulators and neuropeptides implicated in plasticity in amygdala subdivisions following acute and chronic alcohol exposure (e.g. (74)). Additionally, the complexity of the inputs modulating not only BA activity but also the interconnected amygdala subdivisions is far greater than accounted for in this initial model design. Addition of other neural groups, such as the intercalated cells or other projections to the amygdala, such as the prelimbic subdivision of the mPFC, would enable understanding of their interplay with the core modeled here. Thus, the model may provide a means for rapid determination of the putative efficacy of modulators to reduce fear behavior, particularly in the presence of alcohol, thereby identifying novel targets with the highest potential to be further investigated via preclinical in vivo experimentation, potentially streamlining the search for improved treatment targets to ameliorate comorbid stress and alcohol-related mental illnesses.

Altogether, our model demonstrates how plasticity within amygdala circuitry contributes to alcohol’s effects on fear conditioning and make the following predictions: First, the delay in fear extinction in alcohol is mostly induced by greater activation of the BLA after fear acquisition due to alcohol-induced modulation of synaptic weights. Second, the model predicts that both acute and prior repeated alcohol shifts the amygdala network away from the robust extinction regime by inhibiting the activity in the CeA. Third, our model predicts that fear memories formed in acute or after chronic alcohol are more connected to the context, although this prediction is subject to variations in hippocampal activity by alcohol. Finally, our modeling suggests that fear acquisition time may be highly variable and dependent on the specifics of the task, which explains variable results on the influence of alcohol on this time.

## Author Contributions

AL coded and calibrated the model, performed simulations, wrote the text and prepared the manuscript. MLL contributed to interpretation of the results and writing and revising the manuscript. AK led the design of the model and simulations, their interpretation and edited the manuscript.

## References

1. NIAAA. The Cycle of Alcohol Addiction | National Institute on Alcohol Abuse and Alcoholism (NIAAA) [Internet]. 2021 [cited 2023 Dec 4]. Available from: https://www.niaaa.nih.gov/publications/cycle-alcohol-addiction

2. Armstrong JL, Ronzitti S, Hoff RA, Potenza MN. Gender moderates the relationship between stressful life events and psychopathology: Findings from a national study. J Psychiatr Res. 2018 Dec;107:34–41.

3. Moos RH, Moos BS. Treated and Untreated Alcohol-Use Disorders: Course and Predictors of Remission and Relapse. Eval Rev. 2007 Dec;31(6):564–84.

4. Koob GF. Addiction is a Reward Deficit and Stress Surfeit Disorder. Front Psychiatry [Internet]. 2013 [cited 2019 May 21];4. Available from: http://journal.frontiersin.org/article/10.3389/fpsyt.2013.00072/abstract

5. Saladin ME, Brady KT, Dansky BS, Kilpatrick DG. Understanding comorbidity between ptsd and substance use disorders: Two preliminary investigations. Addict Behav. 1995 Sep;20(5):643–55.

6. Ressler RL, Maren S. Synaptic encoding of fear memories in the amygdala. Curr Opin Neurobiol. 2019 Feb;54:54–9.

7. Luchkina NV, Bolshakov VY. Mechanisms of fear learning and extinction: synaptic plasticity– fear memory connection. Psychopharmacology (Berl). 2019 Jan;236(1):163–82.

8. Rescorla RA. Spontaneous Recovery. Learn Mem. 2004 Sep 1;11(5):501–9.

9. Rescorla RA, Heth CD. Reinstatement of fear to an extinguished conditioned stimulus. J Exp Psychol Anim Behav Process. 1975;1(1):88–96.

10. Courtin J, Karalis N, Gonzalez-Campo C, Wurtz H, Herry C. Persistence of amygdala gamma oscillations during extinction learning predicts spontaneous fear recovery. Neurobiol Learn Mem. 2014 Sep;113:82–9.

11. Williams AR, Lattal KM. Rapid reacquisition of contextual fear following extinction in mice: effects of amount of extinction, acute ethanol withdrawal, and ethanol intoxication. Psychopharmacology (Berl). 2019 Jan;236(1):491–506.

12. Likhtik E, Popa D, Apergis-Schoute J, Fidacaro GA, Paré D. Amygdala intercalated neurons are required for expression of fear extinction. Nature. 2008 Jul;454(7204):642–5.

13. Walker DL, Ressler KJ, Lu KT, Davis M. Facilitation of Conditioned Fear Extinction by Systemic Administration or Intra-Amygdala Infusions of d-Cycloserine as Assessed with Fear-Potentiated Startle in Rats. J Neurosci. 2002 Mar 15;22(6):2343–51.

14. Herry C, Ciocchi S, Senn V, Demmou L, Müller C, Lüthi A. Switching on and off fear by distinct neuronal circuits. Nature. 2008 Jul;454(7204):600–6.

15. Pare D, Duvarci S. Amygdala microcircuits mediating fear expression and extinction. Curr Opin Neurobiol. 2012 Aug;22(4):717–23.

16. Williams AR, Lattal KM. Rapid reacquisition of contextual fear following extinction in mice: effects of amount of extinction, acute ethanol withdrawal, and ethanol intoxication. Psychopharmacology (Berl) [Internet]. 2018 Oct 18 [cited 2018 Nov 3]; Available from: http://link.springer.com/10.1007/s00213-018-5057-7

17. Moustafa AA, Gilbertson MW, Orr SP, Herzallah MM, Servatius RJ, Myers CE. A model of amygdala–hippocampal–prefrontal interaction in fear conditioning and extinction in animals. Brain Cogn. 2013 Feb;81(1):29–43.

18. Meyer EM, Long V, Fanselow MS, Spigelman I. Stress Increases Voluntary Alcohol Intake, but Does not Alter Established Drinking Habits in a Rat Model of Posttraumatic Stress Disorder. Alcohol Clin Exp Res. 2013 Apr;37(4):566–74.

19. Logrip ML, Zorrilla EP. Stress history increases alcohol intake in relapse: relation to phosphodiesterase 10A: Stress history alters relapse. Addict Biol. 2012 Sep;17(5):920–33.

20. Dickerson LL, Ferraro DP. Effects of Alcohol on Specific and Environmental Fear. Psychol Rep. 1976 Dec;39(3_suppl):1335–42.

21. Gould TJ. Ethanol disrupts fear conditioning in C57BL/6 mice. J Psychopharmacol (Oxf). 2003 Jan;17(1):77–81.

22. Bisby JA, King JA, Sulpizio V, Degeilh F, Valerie Curran H, Burgess N. Extinction learning is slower, weaker and less context specific after alcohol. Neurobiol Learn Mem. 2015 Nov;125:55–62.

23. Kitaichi K, Minami Y, Amano M, Yamada K, Hasegawa T, Nabeshima T. The attenuation of suppression of motility by triazolam in the conditioned fear stress task is exacerbated by ethanol in mice. Life Sci. 1995;57(8):743–53.

24. Gulick D, Gould TJ. Acute Ethanol Has Biphasic Effects on Short- and Long-Term Memory in Both Foreground and Background Contextual Fear Conditioning in C57BL/6 Mice. Alcohol Clin Exp Res. 2007 Sep;31(9):1528–37.

25. Lattal KM. Effects of ethanol on encoding, consolidation, and expression of extinction following contextual fear conditioning. Behav Neurosci. 2007;121(6):1280–92.

26. Bertotto ME, Bustos SG, Molina VA, Martijena ID. Influence of ethanol withdrawal on fear memory: Effect of D-cycloserine. Neuroscience. 2006 Nov 3;142(4):979–90.

27. Holmes A, Fitzgerald PJ, MacPherson KP, DeBrouse L, Colacicco G, Flynn SM, et al. Chronic alcohol remodels prefrontal neurons and disrupts NMDAR-mediated fear extinction encoding. Nat Neurosci. 2012 Oct;15(10):1359–61.

28. Stephens DN, Ripley TL, Borlikova G, Schubert M, Albrecht D, Hogarth L, et al. Repeated ethanol exposure and withdrawal impairs human fear conditioning and depresses long-term potentiation in rat amygdala and hippocampus. Biol Psychiatry. 2005 Sep 1;58(5):392–400.

29. Quiñones-Laracuente K, Hernández-Rodríguez MY, Bravo-Rivera C, Melendez RI, Quirk GJ. The effect of repeated exposure to ethanol on pre-existing fear memories in rats. Psychopharmacology (Berl). 2015 Oct;232(19):3615–22.

30. Smiley CE, McGonigal JT, Valvano T, Newsom RJ, Otero N, Gass JT. The infralimbic cortex and mGlu5 mediate the effects of chronic intermittent ethanol exposure on fear learning and memory. Psychopharmacology (Berl). 2020 Nov;237(11):3417–33.

31. Janak PH, Tye KM. From circuits to behaviour in the amygdala. Nature. 2015 Jan 15;517(7534):284–92.

32. Ciocchi S, Herry C, Grenier F, Wolff SBE, Letzkus JJ, Vlachos I, et al. Encoding of conditioned fear in central amygdala inhibitory circuits. Nature. 2010 Nov 11;468(7321):277–82.

33. Tovote P, Fadok JP, Lüthi A. Neuronal circuits for fear and anxiety. Nat Rev Neurosci. 2015 Jun;16(6):317–31.

34. LeDoux JE. Emotion circuits in the brain. Annu Rev Neurosci. 2000;23:155–84.

35. Herry C, Ciocchi S, Senn V, Demmou L, Müller C, Lüthi A. Switching on and off fear by distinct neuronal circuits. Nature. 2008 Jul;454(7204):600–6.

36. Milad MR, Quirk GJ. Fear Extinction as a Model for Translational Neuroscience: Ten Years of Progress. Annu Rev Psychol. 2012 Jan 10;63(1):129–51.

37. Pare D, Duvarci S. Amygdala microcircuits mediating fear expression and extinction. Curr Opin Neurobiol. 2012 Aug;22(4):717–23.

38. Duvarci S, Pare D. Amygdala Microcircuits Controlling Learned Fear. Neuron. 2014 Jun;82(5):966–80.

39. Kim J, Pignatelli M, Xu S, Itohara S, Tonegawa S. Antagonistic negative and positive neurons of the basolateral amygdala. Nat Neurosci. 2016 Dec;19(12):1636–46.

40. Zhang X, Kim J, Tonegawa S. Amygdala Reward Neurons Form and Store Fear Extinction Memory [Internet]. bioRxiv; 2019 [cited 2023 Sep 1]. p. 615096. Available from: https://www.biorxiv.org/content/10.1101/615096v1

41. Ehrlich I, Humeau Y, Grenier F, Ciocchi S, Herry C, Lüthi A. Amygdala Inhibitory Circuits and the Control of Fear Memory. Neuron. 2009 Jun 25;62(6):757–71.

42. Krabbe S, Gründemann J, Lüthi A. Amygdala Inhibitory Circuits Regulate Associative Fear Conditioning. Biol Psychiatry. 2018 May;83(10):800–9.

43. Marowsky A, Yanagawa Y, Obata K, Vogt KE. A Specialized Subclass of Interneurons Mediates Dopaminergic Facilitation of Amygdala Function. Neuron. 2005 Dec;48(6):1025–37.

44. Stujenske JM, O’Neill PK, Fernandes-Henriques C, Nahmoud I, Goldburg SR, Singh A, et al. Prelimbic cortex drives discrimination of non-aversion via amygdala somatostatin interneurons. Neuron. 2022 Jul 20;110(14):2258–2267.e11.

45. Wolff SBE, Gründemann J, Tovote P, Krabbe S, Jacobson GA, Müller C, et al. Amygdala interneuron subtypes control fear learning through disinhibition. Nature. 2014 May;509(7501):453–8.

46. Amaya KA, Teboul E, Weiss GL, Antonoudiou P, Maguire J. Basolateral amygdala parvalbumin interneurons coordinate oscillations to drive reward behaviors [Internet]. bioRxiv; 2023 [cited 2023 Aug 10]. p. 2023.08.09.552674. Available from: https://www.biorxiv.org/content/10.1101/2023.08.09.552674v1

47. Mahanty NK, Sah P. Calcium-permeable AMPA receptors mediate long-term potentiation in interneurons in the amygdala. Nature. 1998 Aug 13;394(6694):683–7.

48. Tully K, Li Y, Tsvetkov E, Bolshakov VY. Norepinephrine enables the induction of associative long-term potentiation at thalamo-amygdala synapses. Proc Natl Acad Sci U S A. 2007 Aug 28;104(35):14146–50.

49. Ciocchi S, Herry C, Grenier F, Wolff SBE, Letzkus JJ, Vlachos I, et al. Encoding of conditioned fear in central amygdala inhibitory circuits. Nature. 2010 Nov 11;468(7321):277–82.

50. Cowan JD, Neuman J, van Drongelen W. Wilson–Cowan Equations for Neocortical Dynamics. J Math Neurosci [Internet]. 2016 Dec [cited 2018 Jul 23];6(1). Available from: http://www.mathematical-neuroscience.com/content/6/1/1

51. Carrere M, Alexandre F. A pavlovian model of the amygdala and its influence within the medial temporal lobe. Front Syst Neurosci [Internet]. 2015 Mar 18 [cited 2018 Jul 23];9. Available from: http://www.frontiersin.org/Systems_Neuroscience/10.3389/fnsys.2015.00041/abstract

52. Pape HC, Pare D. Plastic synaptic networks of the amygdala for the acquisition, expression, and extinction of conditioned fear. Physiol Rev. 2010 Apr;90(2):419–63.

53. Blair HT, Schafe GE, Bauer EP, Rodrigues SM, LeDoux JE. Synaptic plasticity in the lateral amygdala: a cellular hypothesis of fear conditioning. Learn Mem Cold Spring Harb N. 2001 Oct;8(5):229–42.

54. Dityatev AE, Bolshakov VY. Amygdala, long-term potentiation, and fear conditioning. Neurosci Rev J Bringing Neurobiol Neurol Psychiatry. 2005 Feb;11(1):75–88.

55. Goosens KA, Maren S. Long-term potentiation as a substrate for memory: evidence from studies of amygdaloid plasticity and Pavlovian fear conditioning. Hippocampus. 2002;12(5):592–9.

56. Kim JJ, Jung MW. Neural circuits and mechanisms involved in Pavlovian fear conditioning: a critical review. Neurosci Biobehav Rev. 2006;30(2):188–202.

57. Maren S. Synaptic mechanisms of associative memory in the amygdala. Neuron. 2005 Sep 15;47(6):783–6.

58. Sigurdsson T, Doyère V, Cain CK, LeDoux JE. Long-term potentiation in the amygdala: a cellular mechanism of fear learning and memory. Neuropharmacology. 2007 Jan;52(1):215–27.

59. Arruda-Carvalho M, Clem RL. Pathway-selective adjustment of prefrontal-amygdala transmission during fear encoding. J Neurosci Off J Soc Neurosci. 2014 Nov 19;34(47):15601– 9.

60. Bloodgood DW, Sugam JA, Holmes A, Kash TL. Fear extinction requires infralimbic cortex projections to the basolateral amygdala. Transl Psychiatry [Internet]. 2018 Dec [cited 2019 May 24];8(1). Available from: http://www.nature.com/articles/s41398-018-0106-x

61. Li SSY, McNally GP. The conditions that promote fear learning: Prediction error and Pavlovian fear conditioning. Neurobiol Learn Mem. 2014 Feb;108:14–21.

62. Prado-Alcalá R, Marina H, Rivas S, Gabriel RR, Quirarte G. Reversal of Extinction by Scopolamine. Physiol Behav. 1994;56(1):27–30.

63. Vlachos I, Herry C, Lüthi A, Aertsen A, Kumar A. Context-Dependent Encoding of Fear and Extinction Memories in a Large-Scale Network Model of the Basal Amygdala. Behrens T, editor. PLoS Comput Biol. 2011 Mar 17;7(3):e1001104.

64. Haubensak W, Kunwar PS, Cai H, Ciocchi S, Wall NR, Ponnusamy R, et al. Genetic dissection of an amygdala microcircuit that gates conditioned fear. Nature. 2010 Nov 11;468(7321):270–6.

65. Vogel E, Krabbe S, Gründemann J, Wamsteeker Cusulin JI, Lüthi A. Projection-Specific Dynamic Regulation of Inhibition in Amygdala Micro-Circuits. Neuron. 2016 Aug;91(3):644– 51.

66. Silberman Y, Shi L, Brunso-Bechtold JK, Weiner JL. Distinct Mechanisms of Ethanol Potentiation of Local and Paracapsular GABAergic Synapses in the Rat Basolateral Amygdala. J Pharmacol Exp Ther. 2007 Oct 5;324(1):251–60.

67. Woodruff AR, Sah P. Inhibition and Synchronization of Basal Amygdala Principal Neuron Spiking by Parvalbumin-Positive Interneurons. J Neurophysiol. 2007 Nov;98(5):2956–61.

68. Roberto M, Madamba SG, Moore SD, Tallent MK, Siggins GR. Ethanol increases GABAergic transmission at both pre- and postsynaptic sites in rat central amygdala neurons. Proc Natl Acad Sci. 2003 Feb 18;100(4):2053–8.

69. Weiner JL, Valenzuela CF. Ethanol modulation of GABAergic transmission: The view from the slice. Pharmacol Ther. 2006 Sep 1;111(3):533–54.

70. Zhu PJ, Lovinger DM. Ethanol potentiates GABAergic synaptic transmission in a postsynaptic neuron/synaptic bouton preparation from basolateral amygdala. J Neurophysiol. 2006 Jul;96(1):433–41.

71. Roberto M, Madamba SG, Stouffer DG, Parsons LH, Siggins GR. Increased GABA release in the central amygdala of ethanol-dependent rats. J Neurosci Off J Soc Neurosci. 2004 Nov 10;24(45):10159–66.

72. Gilpin NW, Herman MA, Roberto M. The Central Amygdala as an Integrative Hub for Anxiety and Alcohol Use Disorders. Biol Psychiatry. 2015 May;77(10):859–69.

73. Läck AK, Diaz MR, Chappell A, DuBois DW, McCool BA. Chronic ethanol and withdrawal differentially modulate pre- and postsynaptic function at glutamatergic synapses in rat basolateral amygdala. J Neurophysiol. 2007 Dec;98(6):3185–96.

74. Roberto M, Gilpin NW, Siggins GR. The Central Amygdala and Alcohol: Role of - Aminobutyric Acid, Glutamate, and Neuropeptides. Cold Spring Harb Perspect Med. 2012 Dec 1;2(12):a012195–a012195.

75. Roberto M, Schweitzer P, Madamba SG, Stouffer DG, Parsons LH, Siggins GR. Acute and chronic ethanol alter glutamatergic transmission in rat central amygdala: an in vitro and in vivo analysis. J Neurosci Off J Soc Neurosci. 2004 Feb 18;24(7):1594–603.

76. Melia KR, Ryabinin AE, Corodimas KP, Wilson MC, LeDoux JE. Hippocampal-dependent learning and experience-dependent activation of the hippocampus are preferentially disrupted by ethanol. Neuroscience. 1996 Jul;74(2):313–22.

77. Ryabinin A, Melia K, Cole M, Bloom F, Wilson M. Alcohol selectively attenuates stress-induced c-fos expression in rat hippocampus. J Neurosci. 1995 Jan 1;15(1):721–30.

78. LaLumiere RT, Buen TV, McGaugh JL. Post-Training Intra-Basolateral Amygdala Infusions of Norepinephrine Enhance Consolidation of Memory for Contextual Fear Conditioning. J Neurosci. 2003 Jul 30;23(17):6754–8.

79. Debiec J, Ledoux JE. Disruption of reconsolidation but not consolidation of auditory fear conditioning by noradrenergic blockade in the amygdala. Neuroscience. 2004;129(2):267–72.

80. Giustino TF, Seemann JR, Acca GM, Goode TD, Fitzgerald PJ, Maren S. β-Adrenoceptor Blockade in the Basolateral Amygdala, But Not the Medial Prefrontal Cortex, Rescues the Immediate Extinction Deficit. Neuropsychopharmacol Off Publ Am Coll Neuropsychopharmacol. 2017 Dec;42(13):2537–44.

81. Skelly MJ, Chappell AM, Ariwodola OJ, Weiner JL. Behavioral and neurophysiological evidence that lateral paracapsular GABAergic synapses in the basolateral amygdala contribute to the acquisition and extinction of fear learning. Neurobiol Learn Mem. 2016 Jan;127:10–6.

82. Silberman Y, Ariwodola OJ, Weiner JL. β1-adrenoceptor activation is required for ethanol enhancement of lateral paracapsular GABAergic synapses in the rat basolateral amygdala. J Pharmacol Exp Ther. 2012 Nov;343(2):451–9.

